# Simulating intervertebral disc cell behavior within 3D multifactorial environments

**DOI:** 10.1101/2019.12.23.886887

**Authors:** L. Baumgartner, J.J. Reagh, M.A. González Ballester, J. Noailly

## Abstract

**Motivation:** Low back pain is responsible for more global disability than any other condition. Its incidence is closely related to intervertebral disc (IVD) failure, which is likely caused by an accumulation of microtrauma within the IVD. Crucial factors in microtrauma development are not entirely known yet, probably because their exploration in vivo or in vitro remains tremendously challenging. In-silico modelling is, therefore, definitively appealing, and shall include approaches to integrate influences of multiple cell stimuli at the microscale. Accordingly, this study introduces a hybrid Agent-based (AB) model in IVD research and exploits network modelling solutions in systems biology to mimic the cellular behavior of Nucleus Pulposus cells exposed to a 3D multifactorial biochemical environment, based on mathematical integrations of existing experimental knowledge. Cellular activity reflected by mRNA expression of Aggrecan, Collagen type I, Collagen type II, MMP-3 and ADAMTS were calculated for inflamed and non-inflamed cells. mRNA expression over long periods of time is additionally determined including cell viability estimations. Model predictions were eventually validated with independent experimental data.

**Results:** As it combines experimental data to simulate cell behavior exposed to a multifactorial environment, the present methodology was able to reproduce cell death within 3 days under glucose deprivation and a 50% decrease in cell viability after 7 days in an acidic environment. Cellular mRNA expression under non-inflamed conditions simulated a quantifiable catabolic shift under an adverse cell environment, and model predictions of mRNA expression of inflamed cells provide new explanation possibilities for unexpected results achieved in experimental research.

## 1 Introduction

Intervertebral discs (IVD) connect the vertebral bodies of the vertebras of the spine. They provide flexibility to the spinal column, act as dampers and transfer the loads from one vertebra to another. Failure of those structures, clinically reflected by degenerative disc disorders or disc herniation, is assumed to be highly related to low back pain, a disorder that causes more global disability than any other (Hoy *et al.*, 2014). The IVD consists of three distinct, specialized tissues, (i) the Nucleus Pulposus (NP), i.e. the gel-like center of the disc, (ii) the Annulus Fibrosus (AF), i.e. a concentrically layered fibrocartilage that laterally confines NP, and (iii) the Cartilage Endplate (CEP) that cranially and caudally separates the NP and the inner AF from the bony endplates of the vertebra (Figure 1).

**Fig. 1.**
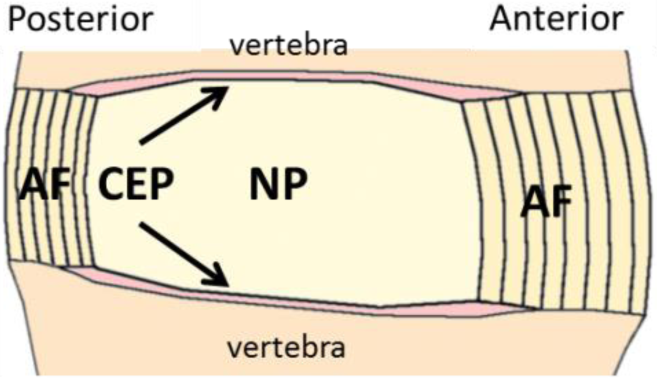
Schematic mid-sagittal cut of a non-degenerated human lumbar intervertebral disc (Adapted from (Noailly, 2009)). Legend: NP: Nucleus Pulposus, AF: Annulus Fibrosus, CEP: Cartilage Endplate.

It is widely accepted that IVD failure is the result of tissue fatigue caused by repetitive (physiological) mechanical loads (e.g. Wade et al. 2016) rather than the result of unique traumatic high impact loads (e.g. Lee & Kwon 2013). A suggested mechanism leading to IVD failure is the accumulation of microtrauma (micro lesions) in the IVD tissue over time, which depends on both, the external loads and the local capacity of the tissue to resist these loads. Yet, such capacity is strongly related to disc cell activity, as cells dynamically build and/or degrade extracellular matrix (ECM). Cell regulation mechanisms are still not fully understood, but local cellular behavior is known to be guided by micro-environmental mechanical and biochemical stimuli, including nutritional stimulation. The latter is of special interest, since IVD are large avascular organs, and local disc cell nutrition relies on solute diffusion (Urban *et al.*, 1977) from the adjacent vertebra. Therefore, important nutritional gradients exist within the NP between regions adjacent to the CEP and the mid-transversal plane (Huang *et al.*, 2014). Biochemical solute diffusion and cell nutrition in the NP is further strain-dependent: external loads influence disc tissue porosity and therefore local amounts of metabolites (Malandrino *et al.*, 2015), which has been called indirect mechanotransduction (Iatridis *et al.*, 2006).

The influence of biochemical stimuli on NP cell activity has been explored through numerous in-vitro studies (e.g. Gilbert et al. 2016; Neidlinger-Wilke et al. 2012; Rinkler et al. 2010; Saggese et al. 2018). While such studies provide valuable information about the overall cell sensitivity to specific microenvironmental cues, they were not designed to capture the spatio-temporal effects of these cues. A further limitation is the limited capacity, mainly because of cost issues, to explore the combined effects of different cell stimuli. Consequently, the cellular behavior over long time periods within a dynamic, multifactorial biochemical environment remain poorly understood. To our best knowledge, no methodology was reported so far that combines findings from different experiments in order to approach, in an integrative way, the complex environmental regulation of IVD cell behavior. Current in-silico modelling approaches to explore the IVD are to a great extent limited to Finite Element modelling (FEM). FEM integrate boundary condition (BC) and structural / composition effects in the local description of disc tissue behaviour, which could be extended to the exploration of both, direct (i.e. the mechanical loads directly sensed by cells on their membranes) and indirect mechanotransduction (Malandrino *et al.*, 2011, 2015; Gu *et al.*, 2014; Van Rijsbergen *et al.*, 2018), even including tissue regulation (Gu, Zhu, and D. 2014; Van Rijsbergen et al. 2018). However, the utility of FEM to simulate the dynamic biological processes responsible for IVD tissue maintenance and failure is limited. In contrast, Agent-based (AB) modelling allows approximating such processes, since it can integrate manifold biological processes, in representative tissue volume elements. AB model studies can realistically improve current understandings of biological or disease mechanisms (Olivares *et al.*, 2016; Holcombe *et al.*, 2012; Santoni *et al.*, 2008) and can be coupled with FEM studies for bottom-up explorations of tissue regulation mechanisms (Ceresa *et al.*, 2018). To the best of our knowledge, no such approach has been developed for the IVD so far. As for other cartilaginous systems, modelling efforts have rather focused on single cell gene regulation (Kerkhofs *et al.*, 2016) or signaling pathways (Melas *et al.*, 2014) instead of collective cell behaviour in representative ECM /tissue.

Consequently, this work focusses on the development and the evaluation of a new methodology to model the behaviour of IVD cells in representative volume elements. The methodology was designed to explicitly and robustly integrate heterogeneous experimental results through AB modelling. Simulations targeted the effect of cell multifactorial biochemical micro-environments to address indirect mechanotransduction phenomena in local tissue volumes, and predicted NP cell behaviour was assessed against individual mRNA expressions and cell viability.

## 2 Methods

The open source software Netlogo version 6.0.2 was used to create an AB model world, consisting of a cube with a volume of 1 mm^3^, represented by 1000 patches. The AB model world was initially seeded with 4000 agents and each agent represented an average NP cell diameter of 10 µm (Hunter *et al.*, 2003). This modelled concentration of 4000 NP cells/ mm^3^ represented the mean cell density found in a human healthy lumbar NP (Maroudas *et al.*, 1975). The biochemical environment of the NP cells was represented as homogenized quantities - to limit the number of agents. Calculation time step-length was set to one hour.

The availability of the biochemical, nutrition related factors lactate (lac) and glucose (glc) were simulated, as potentially important stimuli in indirect mechanotransduction (Huang *et al.*, 2014; Ruiz Wills *et al.*, 2018). Additionally, cellular inflammation was deemed to affect the cellular behavior. The AB model represented inflammation by predicting both, Interleukin 1β (IL-1β) mRNA expression and a corresponding amount of IL-1β protein, since this proinflammatory cytokine appears to be a key factor in IVD degeneration (Vo *et al.*, 2013). To evaluate cellular activity, the respective mRNA expressions of the NP tissue components Aggrecan (Agg), Collagen type II (Col-II) and Collagen type I (Col-I) were simulated, as well as the expression of ECM-degrading enzymes Metalloproteinase 3 (MMP-3) and ADAMTS. Cell activity was differentiated between inflamed and non-inflamed cells (Fig.).

To estimate the effective mRNA expression over time, current cell viability was also calculated, depending on glc, lac and inflammation levels. The overall simulation workflow is described in Figure 3, and the different submodels for inflammation, cell viability and mRNA expression predictions are detailed in the following subsections.

**Fig. 2:**
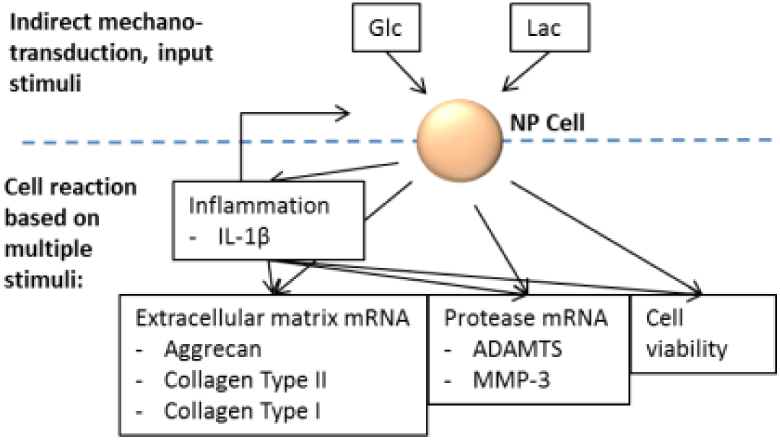
Schematic illustration of included factors in AB model simulations. Glc: glucose, Lac: lactate, IL-1*β*: Interleukin 1*β*

**Fig. 3:**
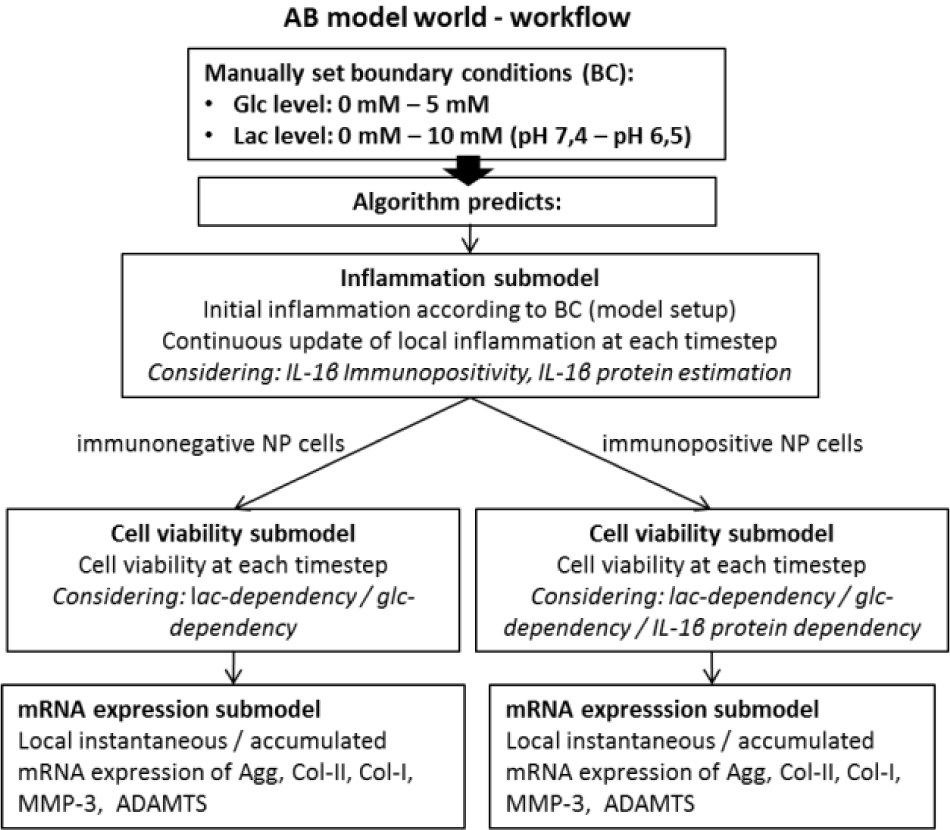
Overview of the AB simulation workflow.

### 2.1 mRNA expression submodel

#### 2.1.1 Prediction of individual mRNA expressions according to varying glc and lac concentrations

To model the influence of a metabolite, i.e. glc or lac (via pH), on cell mRNA expression, continuous mathematical functions were built based on experimental findings, usually reported as discrete semiquantitative measurements (x-fold mRNA expression compared to control). Experimental data were chosen according to cell type (human preferred, followed by bovine), degenerative status of the tissue (not-degenerated preferred over degenerated) and according to the overall consistency of the experimental findings with current knowledge. Measurements associated with both significant findings and tendencies were considered for the development of the mathematical functions. These functions were based on the studies listed in Table 1 and covered the whole physiological spectrum of metabolite concentrations. Each metabolite concentration was related to a specific mRNA expression for each protein. mRNA expressions ranged between 0 and 1, to reflect the lowest and the highest relative levels of expression, respectively (Figure 4).

**Table 1:** Overview of the experimental studies used to estimate mRNA expression based on the solutes glc, lac and the proinflammatory cytokine IL-1β. Deg: degenerated. Non-deg: non-degenerated. Expr: expression

| Study | Cell type | Culture system | Data used for the AB model |  |
| --- | --- | --- | --- | --- |
|  |  |  | Varying micro-environment | mRNA expressions /protein concentrations* |
| Rinkler et al. 2010 | Human, NP, deg and presumably non-deg | Alginate Beads | glc | Agg, Col-I, Col-II, MMP-3 |
| Saggese et al. 2018 | Bovine, NP | Alginate Beads | glc | ADAMTS |
| Moroney et al. 2002 | Porcine, NP | not specified | glc | IL-1β* |
| Gilbert et al. 2016 | Human, NP, deg and non-deg | not specified | pH | IL-1β, Agg, Col-I, MMP-3, ADAMTS |
| Neidlinger-Wilke et al., 2012 | Bovine, NP | Alginate Beads | pH | Col-II |

**Fig. 4:**
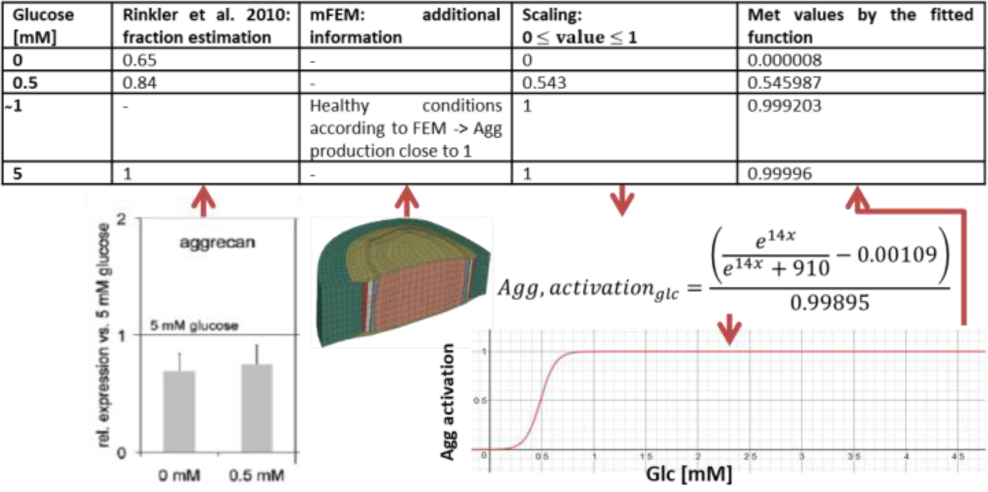
Visualization of the methodology to approximate continuous functions through the example of cellular activity in Agg mRNA expression based on glc concentrations.

The interpolation of discrete experimental measurements suggested a sigmoidal behavior of the change of mRNA expression at critical solute concentrations. Hence, logistic functions were chosen for the interpolation of experimental points, to eventually reflect continuous changes of mRNA expressions. Curve fitting was done by using the free online graphing calculator “Desmos” (https://www.desmos.com/calculator).

To support the function fitting process, an additional datapoint derived from the mechanotransport FEM was included for the glc-dependent mRNA expression curves. Since the mechanotransport simulations predicted glc values as low as ∼1mM in non-deg NP (Ruiz Wills *et al.*, 2018), a datapoint at 1mM glc was added, assuming that cell activity should not be negatively altered at such glc concentration (Figure 4). This point provided robustness for the determination of the start and of the growth rate of the sigmoid. All mRNA expression regulation functions are provided in Table 2.

**Table 2:**
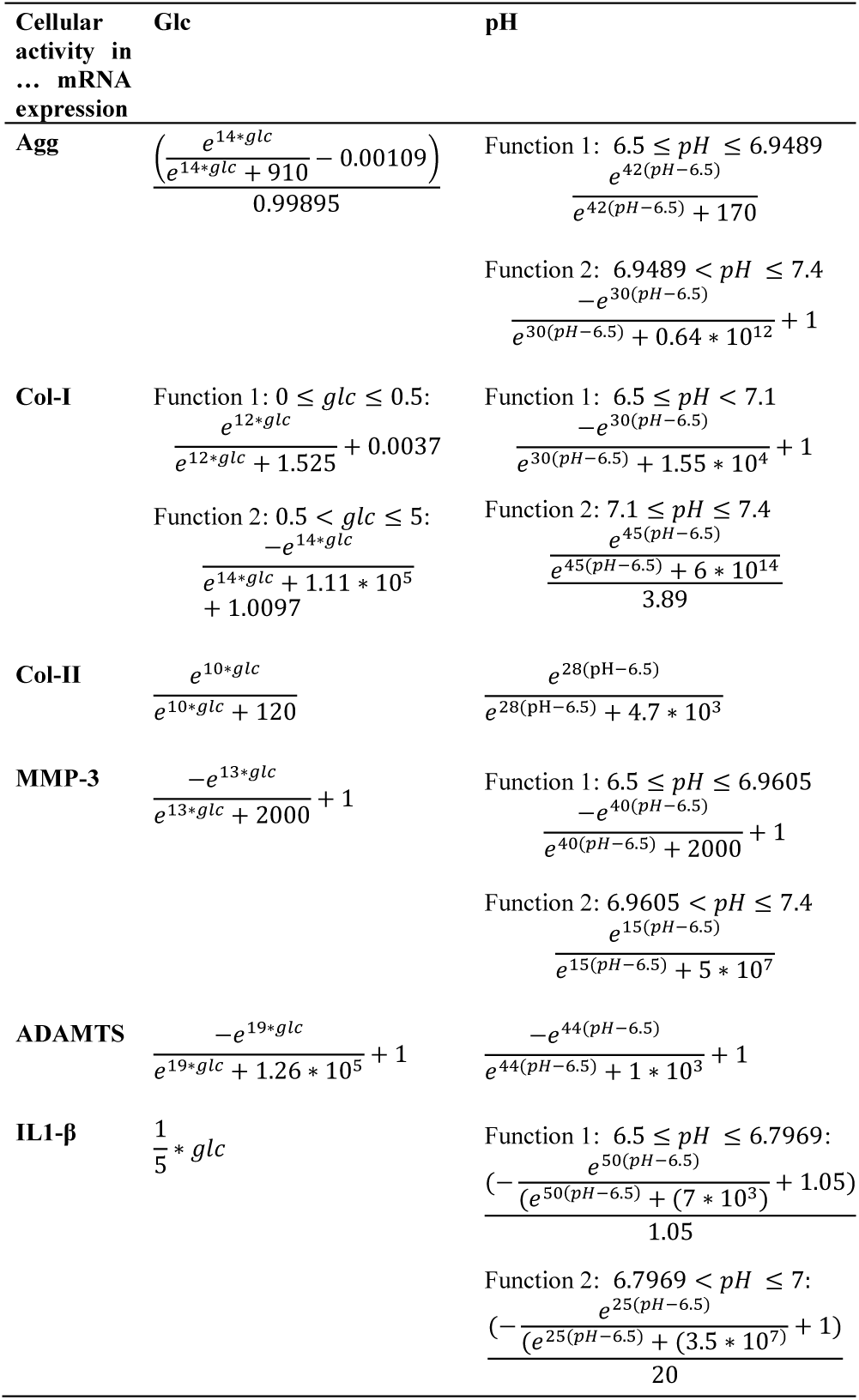
Continuous functions that assign a physiological range of pH and glc, respectively, to a cellular activity in mRNA expression.

#### 2.1.2 Prediction of overall mRNA expression in a multifactorial environment

To combine the respective influences of each stimulus on each mRNA expression, a network-based regulatory model proposed by Mendoza & Xenarios 2006, was used. Briefly, the model equations regulate continuously the nodes of the network (*dx*_*i*_/*dt*) by analytically representing a traditional Boolean integration of the respective effects of activating 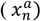 and inhibiting 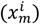 nodes on a nodal input (*ω*_*i*_) (Equation 1). Originally, node regulation also depended on a decay term, which was not relevant in the present work that strictly focused on comparative analyses of the normalized effects of different cell environments. The influence of each inhibiting and/or activating node can be weighted by means of activation (*α*_*n*_) or inhibition (*β*_*n*_) factors. Finally, the way a given nodal input affects the corresponding node is determined by a gain coefficient (*h*).

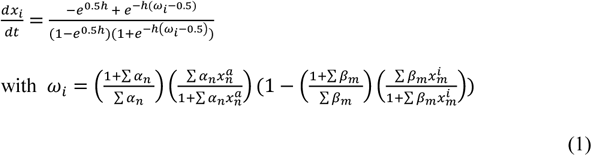

In our regulatory workflow, instead of being nodes regulated by the network, 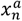 and 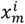 were programmed as dynamical inputs determined by the current metabolic environment of the cells (Figure 5). Accordingly, Equation 1 integrates monofactorial controls of mRNA expressions depending on local glc, lac or IL-1β levels into fully coupled mRNA expressions (*dx*_*i*_/*dt*) within a multifactorial environment

**Fig. 5:**
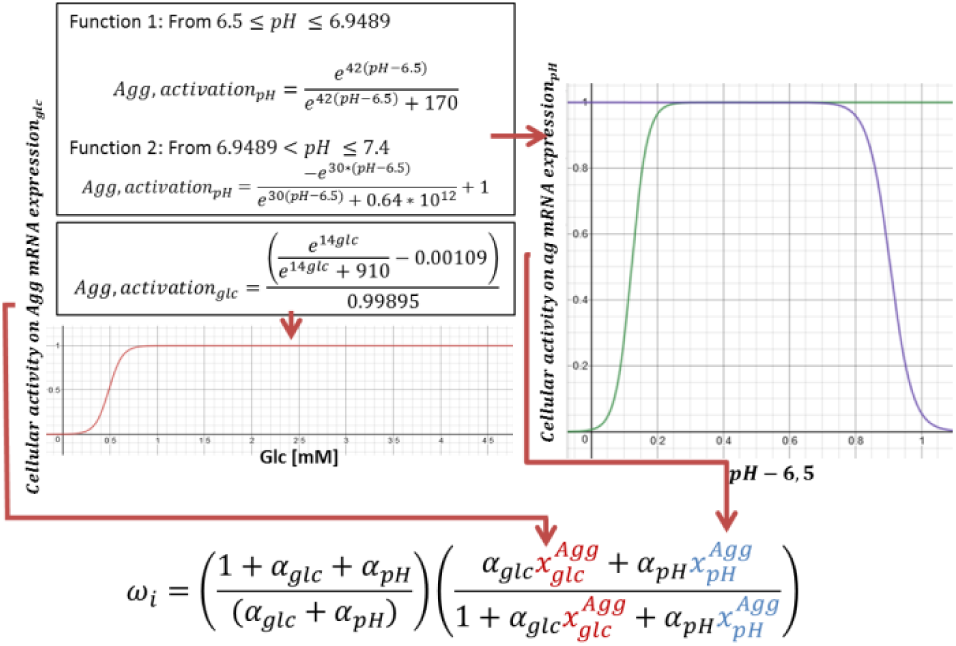
Coupling of experimental results, by using the equation from Mendoza and Xenarios, 2006. Note that pH reaches from 6.5 to 7.4, thus it is represented within the function as values from 0 to 0.9.

The gain factor (h) was generally chosen to linearly relate the input *ω*_*i*_ to the overall activation, and was therefore set to 1. mRNA expressions based on glc and pH were also programmed to have a linear activating effect on *ω*_*i*_, with activation factors (α) of 0.01, according to Mendoza & Xenarios 2006. IL-1β protein was programmed to have an activating effect on MMP-3 and Col-I and to inhibit Agg, Col-II and ADAMTS mRNA expressions, in non-degenerated NP cells according to findings of Le Maitre et al. 2005. The relative activation/inhibition strengths of IL-1β on the mRNA expressions were defined based on these findings. All activation and inhibition factors are summed up in Table 3.

**Table 3.** Overview over estimated weights of stimuli as either activating (α) or inhibiting (β).

| Solute | Activation / inhibition factor | Target-mRNA |
| --- | --- | --- |
| **glc** | $\alpha = 0.01$ | Agg, Col I, Col II, MMP-3, ADAMTS |
| **pH** | $\alpha = 0.01$ | Agg, Col I, Col II, MMP-3, ADAMTS |
| **IL-1β protein** | $\beta = 0.03$ | Agg |
| | $\beta = 0.01$ | Col II |
| | $\alpha = 0.01$ | Col I |
| | $\alpha = 0.05$ | MMP-3 |
| | $\beta = 0.02$ | ADAMTs |

### 2.2 Inflammation submodel

The inflammation submodel estimates the amount of immunopositive cells (cells active in IL-1β mRNA expression) (Figure 6) and a corresponding, amount of IL-1β proteins.

**Fig. 6:**
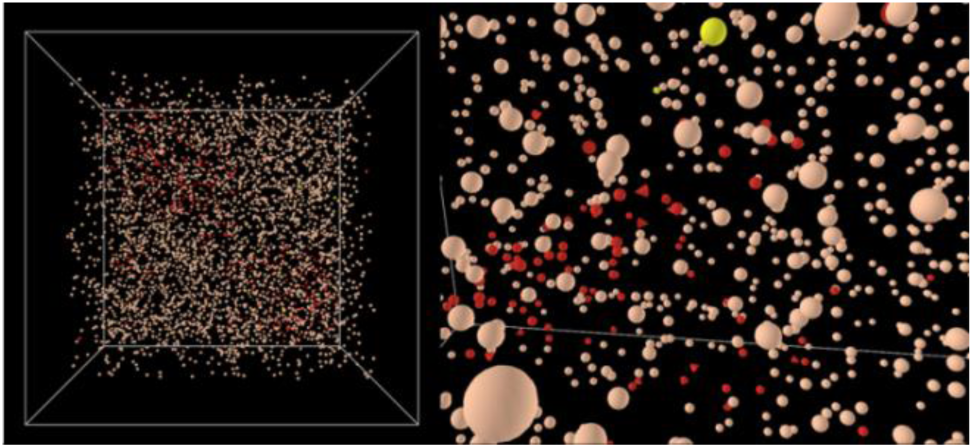
Visualization of immunopositivity within the AB model world (left) and zoom (right). Red: NP-cells immunopositive for IL1-β. Triangle shaped: dead immunopositive cells. Yellow: proliferated cells.

Initial immunopositive NP cell foci were randomly placed within the AB model world and surrounding cell clusters were formed and adapted according to the actual local concentrations of glc, lac and IL-1β protein at each time step (Figure 3). To estimate the quantity of local immunopositive cells, overall IL-1β mRNA expression was determined using the approach from Mendoza & Xenarios 2006. The normalized value of IL-1β mRNA expression was direct proportionally translated into an amount of inflamed cells, assuming a maximum of 20 % of inflamed cells within a non-degenerated human NP (Le Maitre, Freemont and Hoyland, 2005).

Likewise, the regional amount of IL-1β proteins was estimated qualitatively according to the amount of IL-1β mRNA expression by assuming a linear relationship. Half-life of IL-1β proteins was considered by accumulating the relative amount of IL-1β over the current and the previous time step of calculation. To consider the autocrine cell simulation by IL-1β protein (mentioned in e.g. Zou et al. 2017), the accumulated amount of IL-1β protein was normalized to influence the amount of immunopositive cells via feedback loop (Figure 7). Given the low travelling distance of IL-1β due to the size and the short half-life of the protein, the effect of the protein on cell viability and mRNA expression was limited to immunopositive cell-foci. Similar to the solute update submodel, the inflammation submodel was initialized prior to mRNA and cell viability calculations in order to reach a steady state of IL-1β protein concentrations based on user-defined glc and lac concentrations (Figure 3).

**Fig. 7:**
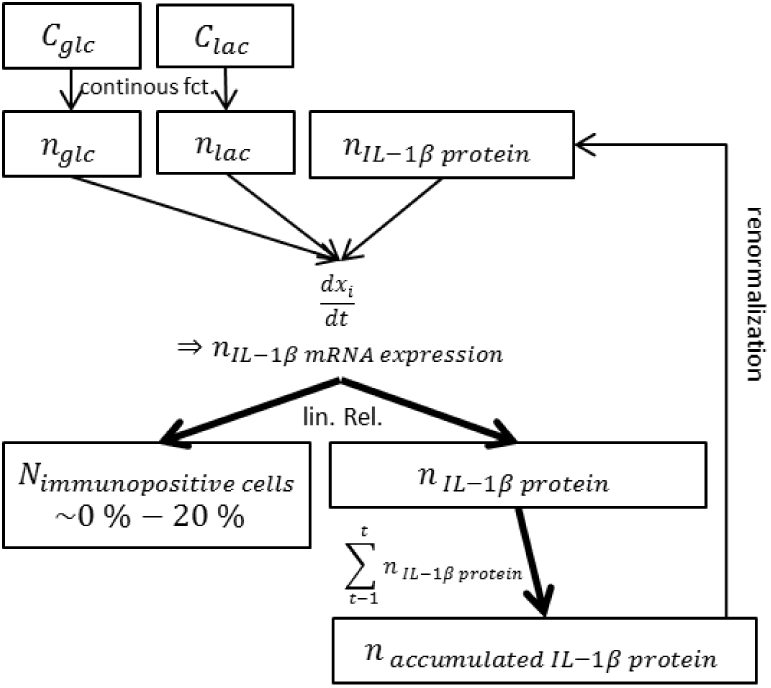
Schematic representation of the inflammation submodel. C: concentration, N: Number, n: normalized

### 2.3 Cell viability submodel

To estimate cell viability, continuous mathematical logistic functions were built (Table 4) according to reported experimental studies (Table 5), reflecting an hourly percentage of decay or proliferation rates based on glc, lac and, in inflamed regions, on IL-1β protein levels. Functions considered both, experimental measurements and important elements of discussions from these studies. E.g. Bibby & Urban 2004 suggested their obtained cell viability to be affected by plenteous cellular glycogen stores due to the high-glucose preculturing media. Consequently, they assumed an adaptation of cells to adverse conditions within the first 24 h of the 48 h experiment. In our model, the mathematical interpretation of this finding consisted in the calculation of a hourly rate of cell death over a time period of 24 h instead of 48 h.

**Table 4:** Continuous functions that determine the percentage of hourly cell-death or cell-proliferation depending on glc and pH levels.

|  | Glc | pH |
| --- | --- | --- |
| **Cell death** | $2.1 * \frac{e^{16*glc}}{e^{16*glc} + 922} - 2.1$ | Function 1: $6.5 \leq pH \leq 6.9$<br>$\frac{1.12 * e^{40(pH-6.5)}}{e^{40(pH-6.5)} + 80} - 1.12$<br>3.57 |
| **Cell proliferation** | - | Function 2: $6.9 < pH \leq 7.4$<br>$\frac{1.12 * e^{42(pH-6.5)}}{e^{42(pH-6.5)} + 5.7 * 10^{10}}$<br>1.775 |

**Table 5:** Overview over experimental studies and their conditions used to estimate cell viability based on the solutes glc, lac and the proinflammatory cytokine IL-1β. AB: Alginate beads. Deg: degenerated. Non-deg: non-degenerated.

| Author | Cell type | Cell culturing | Micro environment |
| --- | --- | --- | --- |
| **Bibby & Urban 2004** | Bovine NP | AB | glc |
| **Gilbert et al. 2016** | Human NP, deg and non-deg | not specified | pH |
| **Shen et al. 2016** | Human NP, non-deg | AB | IL-1β |

Influence of biochemical stimuli on cell viability was programmed to be additive.

Experimentally, cell-death due to IL-1β was determined only for an IL-1β protein concentration of 10 ng/ml (Shen et al. 2016). Because of the incomplete knowledge about physiological concentrations of IL-1β protein within non-degenerated NP and about the critical values of IL-1β protein causing cell death, a threshold of IL-1β protein concentrations leading to cell death was arbitrarily set to half of the maximum IL-1β protein level.

Finally, to estimate the influence of cell viability on mRNA expression over time 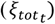, a normalized value of living NP cells (*τ*: current amount, *τ*_0_: initial amount) was multiplied by the instantaneous amount of mRNA expression (*ξ*) at each time step (t). (Equation 9):

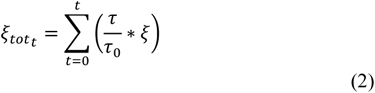

### 2.3 AB model evaluation and validation

#### 2.3.1 mRNA expression submodel evaluation and validation

To qualitatively validate the mRNA prediction, the experimental culture conditions used by Saggese et al. 2018 with regard to pH and glc concentrations were simulated. The authors exposed 3D bovine NP cell cultures to 5.5 mM and 0.55 mM glc at pH 7.0 for 24 h.

5 mM instead of 5.5 mM was chosen for AB model calculations because the selection for solute concentrations is limited to physiological ranges, where 5 mM reflects the highest glc concentration, according to the literature used to build the model regulatory functions (Figure 3/Table 3).

#### 2.3.2 Immunopositivity submodel evaluation

Predictions of the immunopositivity submodel were included in the validation setups for the mRNA expression submodel and the cell viability submodel.

#### 2.3.3 Cell viability submodel validation

To validate cell viability, experimental setups of Horner & Urban 2001 was simulated: during twelve days, cells were exposed to the following conditions: 1) 5 mM glc and at pH 6.0 (for simulations, pH 6.5 was chosen, since this is the lowest value from the physiological range – Figure 3 / Table 2). 2) a glc-free serum, at a pH of 7.4.

Furthermore, the cell microenvironment imposed in the experimental setup of Saggese et al. 2018 (see Section 2.3.1, mRNA expression submodel evaluation and validation) was simulated, to evaluate the influence of cell viability on mRNA expression over time.

## 3 Results

### 3.1 Technical achievements

The model was computationally optimized to simulate the evolution of the system over more than four years in less than 15 min of calculation, with an “ordinary” personal computer (in this study: 16 GB RAM, Intel^(R)^ Core™ i7-7500U CPU @ 2.70GHz (dual core)). More specifically, the AB model could display the normalized IL-1β mRNA expression, the corresponding IL-1β protein concentrations, the current cell viability as well as the instantaneous and accumulated (normalized) mRNA expressions of Agg, Col-I, Col-II, MMP-3 and ADAMTS for inflamed and non-inflamed cells at each timestep (one hour).

### 3.2 mRNA expression submodel

Under glc partial deprivation, (0.55 mM) at pH 7.0 as experimentally imposed by Saggese *et al.*, 2018, the AB model predicts a reduction of the instantaneous mRNA expression of the ECM proteins Agg and Col-II, and an increase of the instantaneous mRNA expression of Col-I, MMP-3 and ADAMTS in the immunonegative cells (Figure 8a).

**Fig. 8a:**
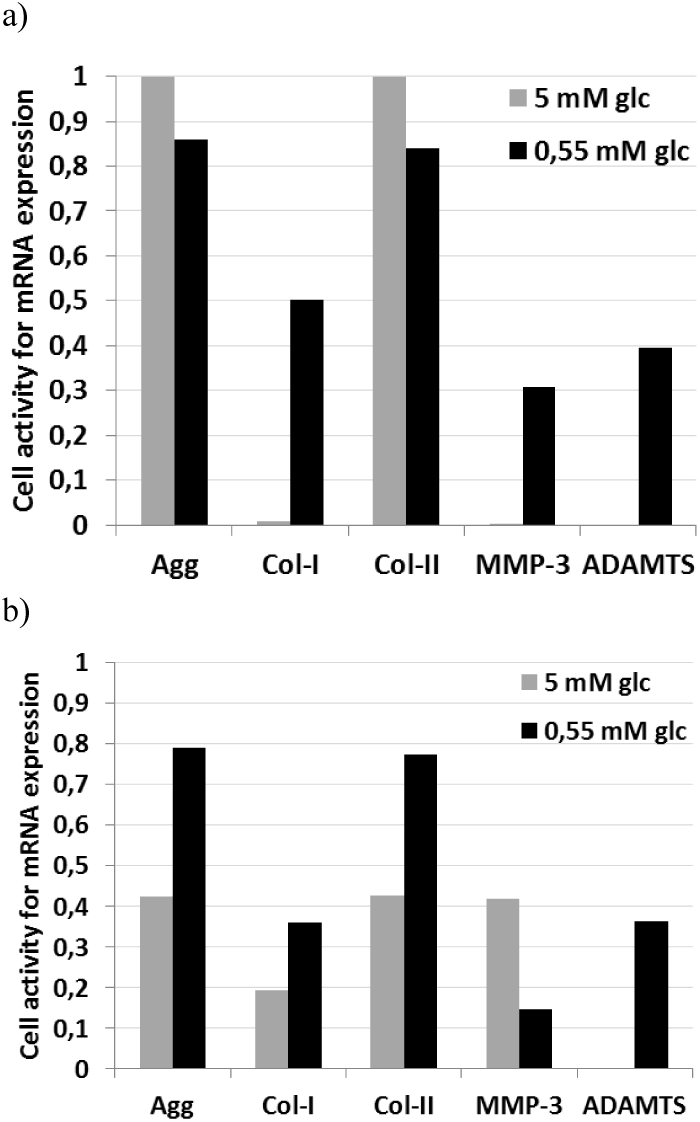
Instantaneous cellular activity in terms of mRNA expression of immunonegative (a) and immunopositive (b) NP cells under glc and pH BC imposed by Saggese et al. 2018 (pH 7.0).

In contrast, simulating the effect of glc partial deprivation at pH 7.0 with immunopositive cells led to increased Agg, Col-I and Col-II mRNA expressions, whereas it reduced MMP-3 mRNA expression (Figure 8b).

### 3.3 Cell viability submodel

Simulations with glc and pH BC similar to those experimentally imposed by Horner & Urban 2001 resulted in:

- 1.2% of cell viability after 2.79 days under glc deprivation.
- cell viability decrease to 71% (78% if immunopositivity is not considered) after three days, to 41% (50% if immunopositivity is not considered) after 7 days and to 11% (14% if immunopositivity is not considered) after 12 days, in acidic environment (pH 6.5) (Figure 9).

Cell decay due to relatively high IL-1β protein levels (i.e. more than half of the maximum permitted) was only activated under acidic conditions.

**Fig. 9:**
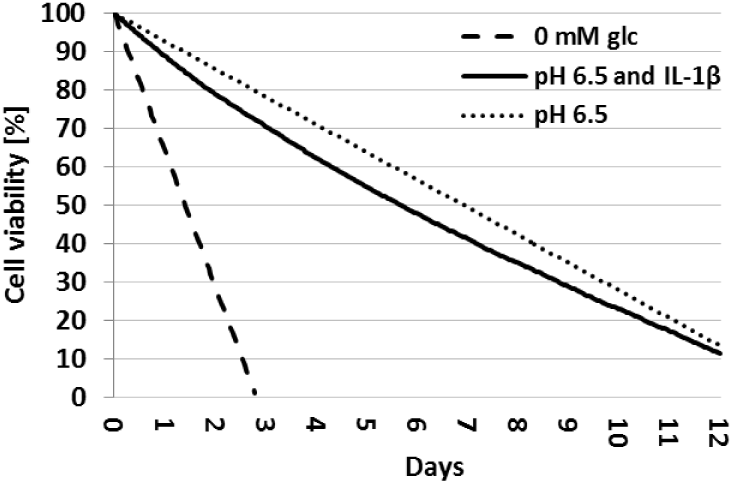
Simulation of cell viability over days in either glc-free medium or acidic environment, with or without IL1 *β* protein.

The influence of cell viability on mRNA expression over time is illustrated in Figure 10 for Agg. mRNA expression slightly increased for non-inflamed cells at 5 mM glc, whereas it decreased in any other condition.

**Fig. 10:**
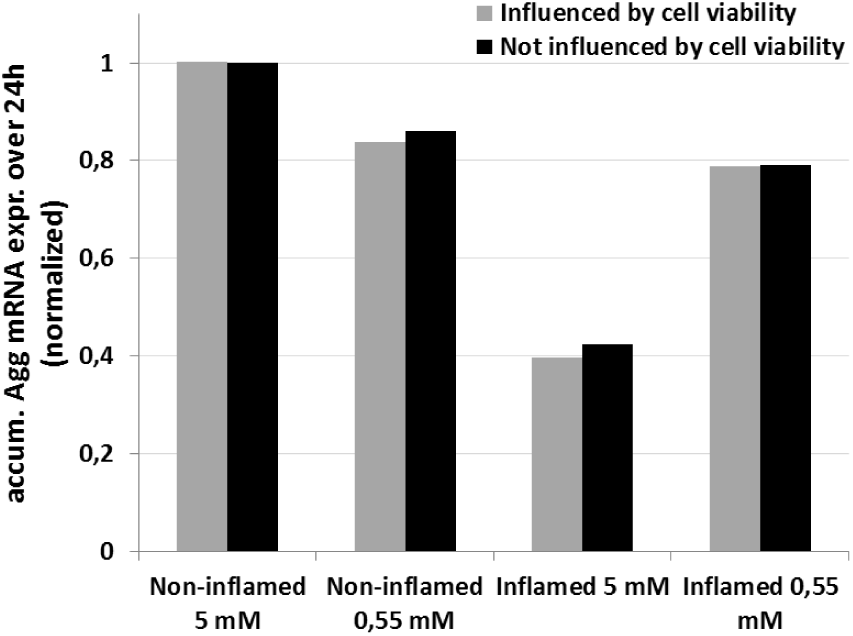
Influence of cell viability on normalized, accumulated (accum.) Agg mRNA expression (expr.). BC as used in Saggese et al. 2018.

## 4 Discussion

To our knowledge, the presented AB model is the first in-silico model that allows predicting the behavior of an intervertebral disc cell population within a multifactorial environment. By its ability to predict cell behavior over long periods of time, it gathers key requirements to reveal cellular dynamics implicated in IVD failure, since microtrauma accumulation is a slow process that we hypothesize to be biologically driven under physiological levels of mechanical loads. Importantly, this approach allows a direct incorporation of experimental data. The experimental data used to build this model were consequently chosen to be upmost consistent with widely accepted assumptions (e.g. a catabolic shift in cell activity due to a decrease of glc concentration). Sometimes contradicting and scarce experimental findings made it difficult to establish general assumptions, e.g. IL-1β mRNA expression was alternatively found to rise (Gilbert et al. 2016) or to fall (Brand et al. 2016) at lower pH levels. In such cases, the measurements associated with the least severe experimental limitations (such as monolayer cultures or limited donors) were used to feed the model. In the following, the performance of the model will be discussed in the light of independent experimental data and current knowledge.

### 4.1 mRNA expression submodel

Continuous functions relating mRNA expressions to glc and lac concentrations are presented in Table 2. In many cases, experimental data suggested more than one sigmoidal shift in mRNA expression over the whole range of physiological concentrations of a stimulating solute (Gilbert *et al.*, 2016). Therefore, a separate function was implemented for each sigmoidal shift (Figure 5).

The fitting of continuous logistic functions generally required mRNA expression measured at three different solute concentrations at least, to obtain (i) a maximum, (ii) a minimum and (iii) an intermediate mRNA expression within a given range of solute concentrations. Hence, the selectable range of solute concentrations within the AB model (Figure 3) was adjusted to the information available through the experimental measurements. In the present study, this is not considered as a limitation, since the used experiments were designed by their respective Authors to cover physiological ranges of glc and lac concentrations. However, if mRNA experimental studies only provide data of mRNA expressions for two concentrations, as it was the case for the relationship between glc and IL1-β (Moroney *et al.*, 2002), it was inevitable to assume a linear relationship between mRNA expression and solute concentrations. Such limitation was primarily due to a current lack of information in the literature. Therefore, valuable information from further experimental research for in-silico approaches such as the current one, shall include mesurements of mRNA expressions at at least three different concentrations of a solute, to duly identify activity shifts. On the one hand, this algorithm is designed to allow straightforward modifications of individual regulatory functions according to new experimental findings. On the other hand, the model can be exploited to identify possible critical cell microenvironments worth to be experimentally explored, e.g. environments around 1 mM glc concentration, pointed out by in-silico calculations of (Ruiz Wills *et al.*, 2018). Furthermore, the algorithm’s structure permits an easy implementation of further factors such as the influence of direct mechanotransduction (magnitude / frequency) on mRNA expression in NP cells (Baumgartner *et al.*, 2019). The effect of oxygen molecule (O_2_) was not considered in this study, since O_2_ levels were reported to have a low effect on the gene expressions hereby simulated, compared to glc and pH (Neidlinger-Wilke *et al.*, 2012). Furthermore, implementation of O_2_ effects is hindered by a lack of experimental data on cell behavior at physiological O_2_ levels (1% - 5% (Mwale *et al.*, 2011)) within the NP.

To build the functions to relate glc or pH, respectively, to mRNA expressions of genes (Table 2), both, significant changes and variation tendencies of mRNA expression were considered among the different experimental reports used. The reason to take into account variation tendencies is, on the one hand, that high variability in cellular behavior comonly seen in experimental studies often hinders to reach significance. On the other hand, we hypothesize that the persistence over time of small changes measured in-vitro can have clear effects over long time periodes. Such hypothesis implies that we hereby consider the chronicity of the cell stimulators studied. Obviously, further model extensions should capture the time variations of these simulators and the effects thereof, including any possible transient effects. At the nutrition level, such variations can be extracted from long-term finite element analyses (Malandrino *et al.*, 2011; Ruiz Wills *et al.*, 2016) and incorporated in the AB model in terms of specific BC history. Coupled to a mechanotransport FEM such as the one presented in (Ruiz Wills *et al.*, 2018), this AB model is able to predict local cellular behavior according to given solute concentrations at the millimeter scale in whatever user-defined region of the NP. As a further development of this AB model, the volume can be subdivided i.e. as ilustrated in Figure 11 as “5 Cubes”. Enriched with a solute-transport and metabolic submodel, the AB model could simulate changes in mRNA expressions because of alterations in solute concentrations at the cell level, i.e. at a higher resolution than the one permited by organ-level FEM. For example, exploration of the influence of microscale tissue consolidations with subsequent inhomogeneous distribution of solutes in the interterritorial matrix would become possible, which might provide further explanation for the – yet unexplained – local cell clustering observed in IVD degeneration (Roberts *et al.*, 2006).

**Fig. 11:**
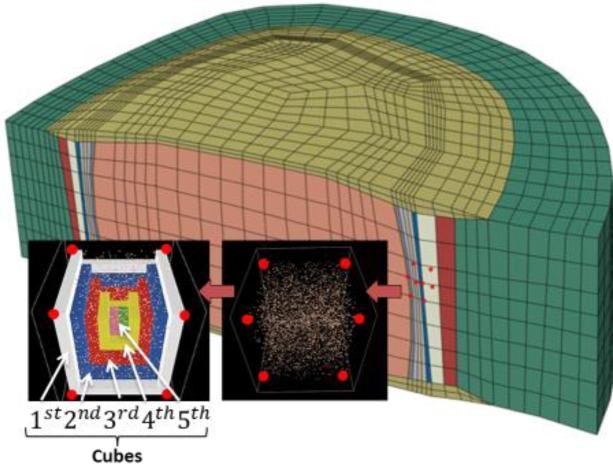
In-house mechanotransport FEM of a human L4-L5 IVD (Ruiz Wills *et al.*, 2018) with coupled AB model within the anterior NP (anterior transition zone). Visualization of NP-cells within this region and subdivision into five different regions (cubes)

The experimental data reported by Saggese et al. 2018 used combinations of pH and glc concentrations (7.0, 0.55 mM) that allowed evaluating the response of the AB model to combined variations of solute concentrations different from those used to build our individual regulatory functions. The predicted behaviour of non-inflamed NP cells to such glc and pH combinations (Figure 8a) is supported by the widely accepted paradigm of catabolic shift in cell activity under glc deprivation, characterized by an overall decline of ECM protein mRNA expression and an increase in protease mRNA expression (Rinkler *et al.*, 2010; Neidlinger-Wilke *et al.*, 2012). Likewise, the predicted augmentation of Col-I expression is well supported by the literature, since a replacement of Col-II with Col-I fibers could be observed in degenerated NPs (Urban and Roberts, 2003; Wuertz *et al.*, 2009).

Remarkably, while the results obtained by Saggese et al. 2018 under glc deprivation support the general assumption that adverse disc cell microenvironments tend to increase Col-I and MMP-3 mRNA expression^1^, they reveal a tendency towards increased Agg and Col-II mRNA expressions, against common expectations. This was not further discussed by the Authors because significance was not reached. Interestingly, our AB model also showed higher Agg and Col-II mRNA expressions under glc deprivation, but only for the immunopositive cells (Figure 10b). Assumingly, this result emerged because IL-1β protein release, and the possible adverse effect thereof on cell activity, is higher at higher glc levels. Hence, less glc, less IL-1β in inflamed foci. This novel insight in cellular behavior within a complex micro-environment suggests that inflammation might shed light on the unexplained experimental findings of Saggese et al. 2018.

Figure 8b shows that at 5 mM glc, the relative mRNA expression of MMP-3 by immunopositive cells was about 0.4, while by immunonegative cells (Figure 8a) kept a minimum amount (i.e. 0.0) of relative MMP-3 mRNA expression. These findings agree with the common association between inflammation and catabolic shift in NP cell activity (Le Maitre *et al.*, 2005). Actually, the catabolic shift in cellular activity was considerable in presence of IL-1β (Figure 8a vs. Figure 8b), such a result is supported by the many evidences that point out sustained inflammation as crucial factor in IVD rupture (e.g. Kepler *et al.*, 2013). However, comparing MMP-3 mRNA expression under glc deprivation for IL-1β immunopositive and immunonegative cells (Figure 8a vs. Figure 8b), model predictions suggest a lower MMP-3 mRNA expression in inflamed cells, even though MMP-3 is programmed to be highly activated by IL-1β (Table 3). This apparent incoherence seems to be related to the use of the equations of Mendoza & Xenarios 2006 (Equation 8) with activation factors that linearly relate a stimulus to the final *ω*. A linear relationship was assumed for the first AB model approach that leads, however, to activation factors below 1. Beta-testing showed that this artefact would vanish with activation factors of *α* ≥ 1, which must be taken into account for future model developments.

To estimate effective inflammation, a linear relationship between IL-1β mRNA expression and the quantity of IL-1β proteins was presumed, which could be roughly observed in experimental studies (Gilbert et al. 2016). However, more detailed knowledge about the relationship between mRNA expression and the corresponding amount of synthesized IL-1β proteins, or about the half-lives of IL-1β protein- or mRNA is required. In the present work, the half-life of IL-1β proteins has been approximated from distantly related studies (Kudo *et al.*, 1990; Larson *et al.*, 2006) and needs to be updated when more accurate data become available.

So far, IL-1β mRNA expression is the only factor that determines the amount of IL-1β protein, but other factors are suggested to affect the levels of IL-1β protein, e.g. MMP-3 might degrade IL-1β (Ito *et al.*, 1996). Thus, further development of the inflammation submodel should include, a more sophisticated feed-back looped network for the dynamic regulation of the effective influence of IL-1β. Furthermore, regulations and effects of additional proinflammatory cytokines are deemed to be crucial in microtrauma development. In particular, the emergence and the role of the proinflammatory cytokine TNF-α, handed as a possible initiating factor in IVD degeneration (Millward-Sadler *et al.*, 2009) should be modelled, but such necessary developments are currently hindered by limited experimental evidences in the literature.

### 4.2 Cell viability submodel

Predicted cell viability is in good agreement with independent, experimental data of Horner & Urban 2001, who observed a decrease in cell viability to about 65% and 50% (cells cultured without oxygen) after three and seven days exposed to low pH. In comparison, our AB model predicted 78% and 50% of viability if the influence of IL1-b is not considered, and 71% and 41% of cell viability otherwise, after three and seven days of exposure to low pH, respectively (Figure 9). Arguably, the minimum pH for our pH-dependent cell regulation curves depended on the lowest pH, i.e. 6.5, found within the experimental studies. In contrast, the experimental study used for independent validation was conducted at pH 6.0. Hence, whether the model might have slightly overestimated the cell viability remains to be explored.

With regard to complete glc deprivation, a cell viability of 1.2% after 2.79 days was predicted by the model (Figure 9), which nicely agrees with the experiments of Horner & Urban 2001 where cells died within three days without glc.

As a limitation, it must be mentioned that the cell viability submodel currently considers cell viability based on additive contributions of biochemical factors (glc, pH and IL-1β protein). Truly, experimental observations (Bibby and Urban, 2004) suggested that coupled factor influences exist. For further model developments, more independent data about cell death in low glc and high acidic environments would be needed to evaluate whether model predictions significantly differ from observed results. Nevertheless, a future development of the cell viability submodel may include a determination of cell viability by using network-based integrative approaches, able to weight the influence of each environmental stimulus individually.

According to the predicted influence of cell viability on the overall mRNA expression over time (24h) (Figure 10), this model allows decoupled quantification of mRNA expression and cell viability to estimate the individual influence of cell death and adverse mRNA expression on the possible subsequent changes in tissue integrity. This is interesting, since cell viability and mRNA expression appear to be decoupled mechanisms (e.g. high glc levels lead to anabolic cell behavior but apparently not to cell proliferation (Bibby and Urban, 2004)). Hence, this model might provide future insight in possibly different pathways of tissue breakdown on different phenotypes of IVD degeneration.

## 5 Conclusion

This investigation aimed to develop and test new modelling and simulation techniques in IVD research for bottom-up virtual exploration of disc tissue regulation and of the negative perturbation thereof. Especially, the presented hybrid model exploited state-of-the-art AB and network modelling approaches. The AB model was fed with experimental data to mimic the cellular behavior in a complex biochemical environment, which stands for a huge challenge, not only in IVD research, but in highly multifactorial disorders in general, where the nature and the control of the physicochemical cellular microenvironments greatly matter. Our new developed methodology successfully integrated discrete, experimental results into continuous cell behavior functions and exploited existing network approaches from systems biology to merge individually weighted stimuli into fully coupled mRNA expressions.

First validation tests showed that the AB model predictions agree with different types of independent experimental findings, and simulations led to novel insights in cellular behavior within a multifactorial environment. Moreover, the AB model provides rational clues to explain for the first time unexpected experimental results retrospectively. This shows the ability of in-silico predictions to contribute to goal-orientated, systematic experimental research and tackle IVD degeneration in an interdisciplinary approach. Thus, feeding a network approach from systems biology with biological data and admitting a direct interference of local environmental conditions seems to be a promising approach for future research.

The strength of this approach is its ability to represent qualitative knowledge /hypotheses when no experimental quantification is available. Furthermore, it enables an integration of heterogeneous, experimental data including cells of different species, differences in cell culturing protocols and degeneration levels of IVD tissue as well as dealing with a lack of data. Importantly, the present methodology enables easy updates of the current regulatory functions provided as soon as new experimental results/findings become available.

Most probably, the NP represents a delicately balanced dynamical system, as local alterations in mRNA expression might lead to local alterations of protein expression. A subsequent, local change in matrix density includes local alterations of tissue porosity at the micrometer scale, which in turn affects local solute concentrations. Therefore, one of the long-term objectives of our work is to couple this AB model with a local FE model, providing continuous feedback about local changes in porosities and the effect of interactions at the microscale level on the millimeter scale tissue integrity, which allows a more holistic understanding and description of the dynamics that finally define the crucial mechanisms for microtrauma accumulation and the great diversity of phenotypes in IVD failure.

## Acknowledgements

The authors thank Dr. Carlos Ruiz Wills for providing the figure of the mechanotransport FEM IVD.

## Funding

This work has been supported by DTIC-UPF, the Whitaker International Fellows and Scholars Program and the Spanish Government (RYC-2015-18888, MDM-2015-0502, HOLOA-DPI2016-80283-C2-1-R).

## Conflict of Interest

none declared.

## Footnotes

1 ADAMTS is excluded from validation as those data were used as input for AB model programing

## References

Baumgartner, L. et al. (2019) Simulation of the Multifactorial Cellular Environment within the Intervertebral disc to better understand Microtrauma Emergence. In, IRC-19-67., pp. 484–485.

Bibby, S.R.S. and Urban, J.P.G. (2004) Effect of nutrient deprivation on the viability of intervertebral disc cells. Eur. Spine J., 13, 695–701.

Brand, F.J. et al. (2016) Acidification changes affect the inflammasome in human nucleus pulposus cells. J. Inflamm., 2, 4–10.

Ceresa, M. et al. (2018) Coupled immunological and biomechanical model of emphysema progression. Front. Physiol., 9, 1–16.

Gilbert, H.T.J. et al. (2016) Acidic pH promotes intervertebral disc degeneration: Acid-sensing ion channel −3 as a potential therapeutic target. Sci. Rep., 6, 1–12.

Gu, W. et al. (2014) Simulation of the Progression of Intervertebral Disc Degeneration due to Decreased Nutrition Supply. Spine (Phila. Pa. 1976)., 39.

Holcombe, M. et al. (2012) Modelling complex biological systems using an agent-based approach. Integr. Biol., 4, 53–64.

Horner, H.A. and Urban, J.P.G. (2001) 2001 Volvo Award Winner in Basic Science Studies : Effect of Nutrient Supply on the Viability of Cells From the Nucleus Pulposus of the Intervertebral Disc. Spine (Phila. Pa. 1976)., 26, 2543–2549.

Hoy, D. et al. (2014) The global burden of low back pain: Estimates from the Global Burden of Disease 2010 study. Ann. Rheum. Dis., 73, 968–974.

Huang, Y.C. et al. (2014) Intervertebral disc regeneration: Do nutrients lead the way? Nat. Rev. Rheumatol., 10, 561–566.

Hunter, C.J. et al. (2003) The three-dimensional architecture of the notochordal nucleus pulposus: novel observations on cell structures in the canine intervertebral disc. J. Anat., 202, 279–291.

Iatridis, J.C. et al. (2006) Effects of Mechanical Loading on intervertebral Disc Metabolism In Vivo (Author Manuscript). J. Bone Jt. Surg. (J Bone Jt. Surg Am), 88, 41–46.

Ito, A. et al. (1996) Communications : Degradation of Interleukin 1 β by Matrix Metalloproteinases Degradation of Interleukin 1 ^N^_L_ by Matrix Metalloproteinases *. 14657–14661.

Kepler, C.K. et al. (2013) The molecular basis of intervertebral disc degeneration. Spine J., 13, 318–330.

Kerkhofs, J. et al. (2016) A qualitative model of the differentiation network in chondrocyte maturation: A holistic view of chondrocyte hypertrophy. PLoS One, 11, 1–27.

Kudo, S. et al. (1990) Clearance and Tissue distribution of Recombinant Human Interleukin 1b in Rats. Cancer Res. (Cancer Res), 50, 5751–5755.

Larson, J.W. et al. (2006) Biologic modification of animal models of intervertebral disc degeneration. J. Bone Joint Surg. Am., 88 Suppl 2, 83–87.

Lee, H.-W. and Kwon, Y.-M. (2013) Traumatic Intradural Lumbar Disc Herniation without Bone Injury. Korean J. Spine, 10, 181–184.

Le Maitre, C.L. et al. (2005) The role of interleukin-1 in the pathogenesis of human intervertebral disc degeneration. Arthritis Res. Ther., 7, R732–R745.

Malandrino, A. et al. (2015) Poroelastic modeling of the intervertebral disc: A path toward integrated studies of tissue biophysics and organ degeneration. MRS Bull., 40, 324–332.

Malandrino, A. et al. (2011) The effect of sustained compression on oxygen metabolic transport in the intervertebral disc decreases with degenerative changes. PLoS Comput. Biol., 7.

Maroudas, A. et al. (1975) Factors involved in the nutrition of the human lumbar intervertebral disc: cellularity and diffusion of glucose in vitro. J. Anat. (J. Anat.), 120, 113–130.

Melas, I.N. et al. (2014) Modeling of signaling pathways in chondrocytes based on phosphoproteomic and cytokine release data. Osteoarthr. Cartil., 22, 509–518.

Mendoza, L. and Xenarios, I. (2006) A method for the generation of standardized qualitative dynamical systems of regulatory networks. Theor. Biol. Med. Model. (Theor Biol Med Model., 3, 13.

Millward-Sadler, S.J. et al. (2009) Regulation of catabolic gene expression in normal and degenerate human intervertebral disc cells: implications for the pathogenesis of intervertebral disc degeneration. Arthritis Res. Ther., 11, R65.

Moroney, P.J. et al. (2002) PH and anti-inflammatory agents modulate nucleus pulposus cytokine secretion. In, Proceedings of the NASS 17th Annual Meeting / The Spine Journal., p. 47S–128S.

Mwale, F. et al. (2011) Effect of oxygen levels on proteoglycan synthesis by intervertebral disc cells. Spine (Phila. Pa. 1976)., 36, 131–138.

Neidlinger-Wilke, C. et al. (2012) Interactions of environmental conditions and mechanical loads have influence on matrix turnover by nucleus pulposus cells. J. Orthop. Res. (J Orthop Res), 30, 112–121.

Noailly, J. (2009) Model developments for in silico studies of the lumbar spine biomechanics - Tesi doctoral. UPC Departament de Ciència dels Materials i Enginyeria Metallurgica, pp. 1–111.

Olivares, A.L. et al. (2016) Virtual exploration of early stage atherosclerosis. Bioinformatics, 32, 3798–3806.

Van Rijsbergen, M. et al. (2018) Comparison of patient-specific computational models vs. clinical follow-up, for adjacent segment disc degeneration and bone remodelling after spinal fusion. PLoS One, 13, 1–24.

Rinkler, C. et al. (2010) Influence of low glucose supply on the regulation of gene expression by nucleus pulposus cells and their responsiveness to mechanical loading. J. Neurosurg. Spine J Neurosurg Spine, 13, 535–542.

Roberts, S. et al. (2006) Histology and pathology of the human intervertebral disc. J. Bone Jt. Surg. - Ser. A, 88, 10–14.

Ruiz Wills, C. et al. (2016) Simulating the sensitivity of cell nutritive environment to composition changes within the intervertebral disc. J. Mech. Phys. Solids, 90, 108–123.

Ruiz Wills, C. et al. (2018) Theoretical Explorations Generate New Hypotheses About the Role of the Cartilage Endplate in Early Intervertebral Disk Degeneration. Front. Physiol., 9, 1–12.

Saggese, T. et al. (2018) Differential Response of Bovine Mature Nucleus Pulposus and Notochordal Cells to Hydrostatic Pressure and Glucose Restriction. Cartilage, 0, 1–13.

Santoni, D. et al. (2008) Implementation of a regulatory gene network to simulate the TH1/2 differentiation in an agent-based model of hypersensitivity reactions. Bioinformatics, 24, 1374–1380.

Shen, J. et al. (2016) SIRT1 Inhibits the Catabolic Effect of IL-1 beta Through TLR2/SIRT1/NF-kappa B Pathway in Human Degenerative Nucleus Pulposus Cells. Pain Physician, 19, E215–E226.

Urban, J.P. et al. (1977) Nutrition of the intervertebral disk. An in vitro study of solute transport. Clin Orthop Relat Res, 129, 101–14.

Urban, J.P.G. and Roberts, S. (2003) Degeneration of the intervertebral disc. Arthritis Res. Ther. (Arthritis Res Ther), 5, 120–130.

Vo, N. V. et al. (2013) Expression and regulation of metalloproteinases and their inhibitors in interertebral disc aging and degeneration. 13, 331–341.

Wade, K.R. et al. (2016) ISSLS Prize Winner: Vibration Really Disrupt the Disc: A Microanatomical Investigation. Spine (Phila. Pa. 1976)., 41, 1185–98.

Wuertz, K. et al. (2009) In vivo remodeling of intervertebral discs in response to short- and long-term dynamic compression. J. Orthop. Res. (J Orthop Res), 27, 1235–1242.

Zou, J. et al. (2017) Effect of a low-frequency pulsed electromagnetic field on expression and secretion of IL-1β and TNF-α in nucleus pulposus cells. J. Int. Med. Res., 45, 462–470.

